# A target-specific assay for rapid and quantitative detection of Mycobacterium chimaera DNA in environmental and clinical specimens

**DOI:** 10.1101/105213

**Authors:** Enrique Zozaya-Valdés, Jessica L. Porter, John Coventry, Janet A. M. Fyfe, Glen P. Carter, Anders Gonçalves da Silva, Torsten Seemann, Paul D. R. Johnson, Andrew James Stewardson, Ivan Bastian, Sally A. Roberts, Benjamin P. Howden, Deborah A. Williamson, Timothy P. Stinear

**Affiliations:** Department of Microbiology and Immunology, The Doherty Institute for Infection and Immunity, University of Melbourne, Victoria, Australia; Microbiological Diagnostic Unit Public Health Laboratory, Department of Microbiology and Immunology, The Doherty Institute for Infection and Immunity, University of Melbourne, Victoria, Australia; Victorian Infectious Diseases Laboratory,The Doherty Institute for Infection and Immunity, Victoria, Australia; Victorian Life Sciences Computational Initiative, University of Melbourne, Victoria, Australia; Austin Health; SA Pathology, South Australia, Australia; Mycobacterial Laboratory, LabPlus, Auckland, New Zealand

**Author notes:** Address correspondence to Timothy P. Stinear.

## Abstract

*Mycobacterium chimaera* is an opportunistic environmental mycobacterium, belonging to the *Mycobacterium intracellulare* complex. Although most commonly associated with pulmonary disease, there has been growing awareness of invasive *M. chimaera* infections following cardiac surgery. Investigations suggest world-wide spread of a specific *M. chimaera* clone, associated with contaminated hospital heater-cooler units used during the surgery. Given the global dissemination of this clone, its potential to cause invasive disease, and the laboriousness of current culture-based diagnostic methods, there is a pressing need to develop rapid and accurate diagnostic assays, specific for *M. chimaera*. Here, we assessed 354 mycobacterial genome sequences and confirmed that *M. chimaera* is a phylogenetically coherent group. *In silico* comparisons indicated six DNA regions present only in *M. chimaera*. We targeted one of these regions and developed a TaqMan qPCR assay for *M. chimaera* with a detection limit of 10 CFU in whole blood. *In vitro* screening against DNA extracted from 40 other mycobacteria and 22 bacterial species from 21 diverse genera confirmed *in silico* predicted specificity for *M. chimaera*. Screening 33 water samples from heater cooler units with this assay highlighted the increased sensitivity of PCR compared to culture, with 15 of 23 culture negative samples positive by *M. chimaera* qPCR. We have thus developed a robust molecular assay that can be readily and rapidly deployed to screen clinical and environmental specimens for *M. chimaera*.

## Introduction

*Mycobacterium chimaera* is an environmental mycobacterium and infrequent pathogen, most commonly linked with pulmonary disease (1-8). Interest in *M. chimaera* has heightened with global reports of invasive infections (including endocarditis and vascular graft infections associated with the use of LivaNova PLC (formerly Sorin Group Deutschland GmbH) Stöckert 3T heater-cooler units during cardiac surgery. The most plausible hypothesis for this widespread contamination is a point-source outbreak, although the underlying causative factors are not currently known (9-16). Phylogenetic comparisons of 16S–23S rRNA internal transcribed spacer (ITS) sequences, and/or partial *rpoB* or *hsp65* sequences (2, 5-7, 17, 18) suggest *M. chimaera* as a distinct entity within the *M. intracelluare* complex (6) and two recent population genomic analyses have confirmed this relationship (8, 13). The complete 6,593,403 bp genome sequence of *M. chimaera* ANZ045 revealed a single circular 6,078,672 bp chromosome and five circular plasmids ranging in size from 21,123 bp to 324,321 (8). *M. chimaera* is slow-growing, therefore current culture-based laboratory methods, followed by Sanger sequencing of amplicons for one or more combinations of conserved sequence regions, or line-probe hybridization assays are not amenable to timely and specific detection of this pathogen. This delay carries significant clinical, health provision, and medico-legal implications as patients may be exposed to contaminated machines during this turn-around time of up to 6-8 weeks. A rapid and reliable diagnostic tool is urgently needed to support clinical management of patients and to establish the efficacy of heater-cooler unit decontamination procedures. Here we addressed this issue by using comparative genomics to identify DNA sequences present in *M. chimaera* and absent from other mycobacteria. We describe the initial development and validation of a sensitive, specific and quantitative PCR assay for identification of *M. chimaera* in both clinical and environmental samples.

## Materials and methods

### Bacterial strains and genome sequences

*Mycobacterium chimaera* strain DMG1600125 (a 2016 HCU isolate from New Zealand) was used for spiking experiments (8). The mycobacterial genome sequences used in this study are listed in Table S1. *M. chimaera* was grown on Brown and Buckle whole-egg media, Middlebrook 7H9 broth or Middlebrook 7H10 agar (Becton Dickinson) supplemented with 10% (v/v) oleic acid albumin dextrose complex (OADC; Difco) or Middlebrook 7H10 agar. Cultures were incubated without shaking at 37°C. *M. chimaera* colony counts were obtained by spotting 3 µL microliter volumes of six, 10-fold serial dilutions of a *M. chimaera* culture suspensions in quintuplicate on two Middlebrook 7H10 agar plates. The colonies were counted after incubation for four weeks at 37°C.

### Genomic DNA extraction methods, M. chimaera culture and environmental isolation

Purified *M. chimaera* genomic DNA for TaqMan assay validation was extracted from 50 mg wet weight cell pellets as described (19) and measured by fluorimetry using the Qubit and the *High Sensitivity* DNA kit (Thermofisher). For spiking experiments in blood, *M. chimaera* DNA was extracted from 100 μL volumes of whole blood, using the Qiagen Blood & Tissue DNA extraction kit. Purified DNA was eluted from the columns in a 200 μL volume of 10 mM Tris (pH 8.0) (Qiagen). Total bacteria were concentrated from 30-1000 mL volumes of water collected from heater-cooler units by filtration through 47 mm, 0.22 μM mixed cellulose ester Millipore membranes. Immediately after filtration, membranes were aseptically placed in sterile 50 ml plastic tubes and stored at −70° C. DNA was extracted from membrane concentrate using the MoBio PowerWater DNA isolation kit following the manufacturer’s instructions (MoBio) with an additional physical disruption step consisting of 2 × 20 sec at 5000 rpm in a Precellys 24 tissue homogenizer. To prevent cross contamination, a sterilized filtration device was used for each sample and sterile, distilled water extraction blanks were filtered and processed (100 mL volumes) at a frequency of one for every 10 test samples. Culture isolation of *M. chimaera* from 50 mL volumes of water samples was undertaken as described (20).

### Population structure and phylogenetic analysis

Snippy v3.1 (https://github.com/tseemann/snippy) was used to align Illumina sequence read data or *de novo* assembled contigs from *M. chimaera* and related mycobacterial genomes against the fully-assembled, complete MC_ANZ045 reference genome to call core genome single nucleotide polymorphism (SNP) differences and generate pairwise sequence alignments. Hierarchical Bayesian clustering (hierBAPS) was performed using these core whole-genome SNP alignments as input to assess population structure (a prior of 6 depth levels and a maximum of 20 clusters was specified) (21), with phylogenies inferred using FastTree v2.1.8 under a GTR model of nucleotide substitution (22). Pairwise SNP analysis between groups of genomes was performed using a custom R script (https://github.com/MDU-PHL/pairwise_snp_differences). Recombination detection was performed using ClonalFrameML v1.7 (23).

### In silico subtractive hybridization and target identification

To identify regions of DNA present in *M. chimaera* but absent from other mycobacteria, the Illumina sequence reads of 46 *M. chimaera* isolates from Australia and New Zealand and eight publicly available *M. intracellulare* genomes (Table S1) were aligned using BWA MEM v0.7.15-r1140 (https://arxiv.org/abs/1303.3997) to a complete *M. chimaera* reference genome (MC_ANZ045) (8). The read depth at each position was examined to identify those positions in the reference genome that were present across all *M. chimaera* isolates but absent from all *M. intracellulare* genomes. These genomic regions were extracted from MC_ANZ045 and compared against the NCBI Genbank non-redundant (*nt*) nucleotide database using NCBI BLAST v2.5.0 (CITE?) with parameters -*remote -max*_*target*_*seqs 100 -task blastn -outfmt* “*6 std qcovs staxid ssciname*”. Resulting BLAST hits that were missing a taxon name were retrieved from the NCBI taxonomy database using the taxon id. Ignoring BLAST hits against bonafide *M. chimaera* sequences, the query alignment positions for every hit were extracted and were used to obtain all the sequence segments that had no hits against the Genbank *ntxs* database and that were greater than 500 bp in length. For this, *bedtools complement* and *getfasta* tools were used (24). The sequence segments thus obtained were considered candidate *M. chimaera*-specific genomic regions. The presence of these regions across a wider collection of *M. chimaera* was assessed by downloading all *M. chimaera* genome sequence reads present in the NCBI sequence read archive SRA as of October 2016 (Table S1) and processing through the Nullarbor pipeline v1.2 (https://github.com/tseemann/nullarbor). The output information was used to filter out poor quality or non-*M. chimaera* reads sets based on G+C content significantly below 66%, an average read depth below 30, a total contig length above 8Mb, predicted rRNA genes greater than four, or a total of sequence aligned to the reference genome below 70% (Table S1). Using Snippy again, all M. chimaera genomes identified above were mapped to a version of the MC_ANZ045 reference genome in which the non*-M. chimaera*-specific sequence regions had been hard masked. The resulting multiple sequence alignment was parsed using a custom Perl script to identify those *M. chimaera*-specific regions that were present in all *M. chimaera* genomes. These DNA sequences were inspected further for development of *M. chimaera* TaqMan PCR diagnostic assays. TaqMan primers and probes (Sigma Oligonucleotides) were designed using Primer3 (25) and Primer-BLAST against NCBI *nt* database was used to check that the primers and probes designed were specific to *M. chimaera*. TaqMan probe 1970-P were labeled with the fluorescent dye 6-carboxyfluorescein (FAM) at the 5′ end and a nonfluorescent quencher at the 3′ end (Sigma Oligonucleotides). To assess the context of these *M. chimaera*-specific regions, AlienHunter v1.4 was used to screen the MC_ANZ045 genome for DNA compositional bias, indicative of horizontally acquired DNA (26).

### TaqMan quantitative PCR

TaqMan PCR mixtures contained 2 μl of template DNA, 0.4-μM concentrations of each primer, a 0.2 μM concentration of the probe, SensiFAST Probe Lo-ROX (1x) mix (Bioline), and TaqMan exogenous internal positive control (IPC) reagents (Applied Biosystems) in a total volume of 20 µl. Amplification and detection were performed with the Mx3005P (Stratagene) using the following program: 40 cycles of 95°C for 10 s and 60°C for 20 s. DNA extracts were tested in at least duplicate, and negative and positive template controls were included in each run. Standard curves were prepared using eight, 10-fold serial dilutions of M. chimaera genomic DNA at an initial concentration of 120 ng/µL, tested in triplicate. The percentage PCR amplification efficiency (E) for the TaqMan assay was calculated from the slope (C) of the standard curve E = (10^(-1/C))*100. Cycle threshold values for unknown samples were converted to genome equivalents by interpolation, with reference to the standard curve of Ct versus dilutions of known concentrations of M. chimaera genomic DNA. The mass in femtograms of a single M. chimaera genome was estimated as 6.59 fg, using the formula M = (N)*(1.096e-21), where M = mass of the single double-stranded M. chimaera NZ045 reference genome and N = 6593403, which is the length of the M. chimaera NZ045 reference genome, and assuming the average MW of a double-stranded DNA molecule is 660 g/mol. Analyses were performed using Graphpad Prism v6.0h.

## Results

### *Assessment of* M. chimaera *population structure*

To identify DNA segments present only in *M. chimaera* genomes we first assessed the phylogenetic coherence of “*M. chimaera*” as a species. Using 96 mycobacterial genome sequences, comprising 63 *M. chimaera* genomes from North America, Australia, and New Zealand, and 33 other related, publicly available mycobacteria from the *Mycobacterium avium-intracelluare* complex, we conducted whole genome pairwise comparisons of the 96 taxa to the *M. chimaera* ANZ045 complete reference chromosome. The 63 *M. chimaera* genomes included 49 HCU-associated and 14 previously described patient isolates, not all of which were associated with Stöckert 3T HCU contamination (8, 12). These comparisons identified 448,878 variable nucleotide positions in a 2,340,885 bp core genome. A robust phylogeny inferred from the alignments strongly suggested that *M. chimaera* forms a monophyletic lineage within the *M. intracellulare* complex (Fig. 1A) (8, 13). Bayesian analysis of population structure (BAPS) using these same data confirmed this clustering (Fig. 1A). Interestingly, this assessment indicated that a publicly available isolate originally identified as *Mycobacterium intracellulare* (strain MIN_052511_1280) was in fact *M. chimaera*. The mean number of SNPs between any pair of the 63 *M. chimaera* isolates, and MIN_052511_1280 (BAPS-3), not adjusted for recombination, was 115 SNPs (range: 1 - 3,024 and IQR: 13 - 31), highlighting restricted core genome variation within this species, particularly given the large 6.5 Mb genome size. In comparison, the mean number of SNPs between 15 *M. intracellulare*-complex genomes (BAPS-2) was 24,134 SNPs (range: 13 - 39,109 and IQR: 14,780 - 33,138) (Fig. 1A). We then extended this analysis to assess an additional 257 publicly available *M. chimaera* and related mycobacterial genome sequences (Table S1). Pairwise whole genome comparisons of this larger data set were performed against the ANZ045 reference genome. Population structure analysis indicated 303 mycobacterial genomes fell within BAPS-3 (Table S1). Pairwise comparisons were again performed against the ANZ045 using only these 303 genomes, with five other genome sequences from BAPS-2 included for context. The alignment was filtered to remove sites from the alignment affected by recombination and a phylogeny was inferred from the resulting 10,166 variable nucleotide positions (Fig. S1. Fig. 2). The mean number of SNPs between the 303 genomes was 268 (range: 0 - 3,211 and IQR: 9 - 62). This analysis confirmed that *M. chimaera* does indeed form a monophyletic lineage, providing a robust genetic definition for the species. As previously reported, HCU-associated isolates from around the world formed a distinct sub-clade within this lineage (Fig. 2) (8, 13).

**Fig. 1.**
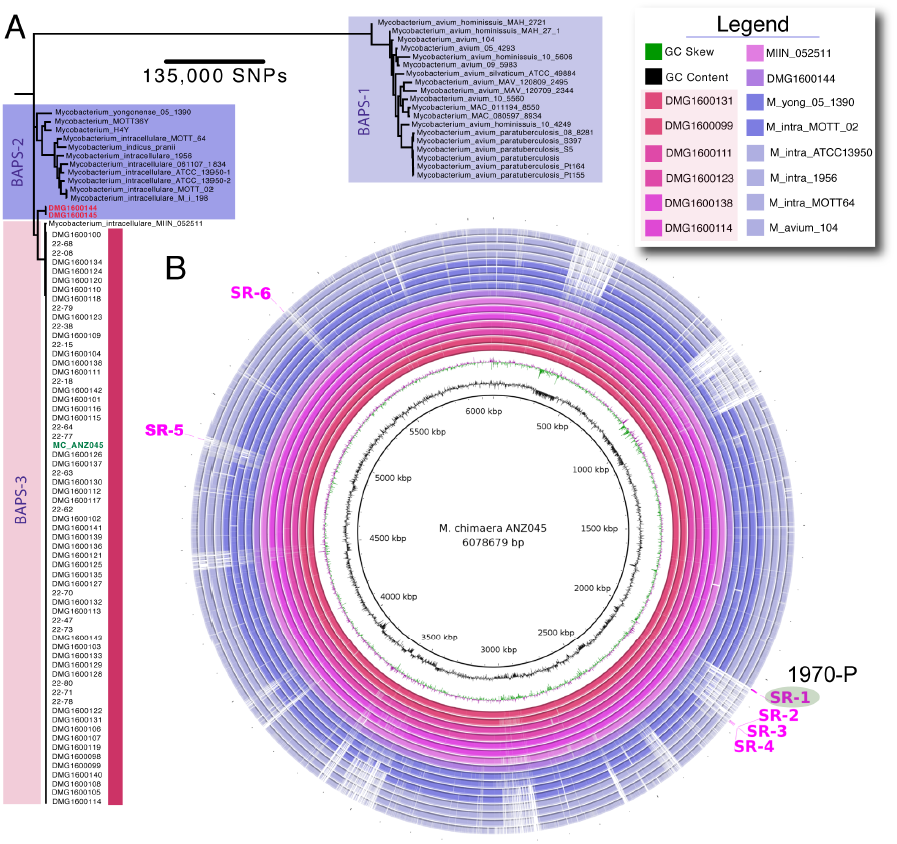
**Phylogenetic and population structure analysis of *M. chimaera*** (A): Core genome maximumlikelihood phylogeny of 63 *M. chimaera* and 33 other, related mycobacteria, based on alignment of 448,878 variable nucleotide positions. The tree was inferred with FastTree using a GTR model of nucleotide substitution. All major branches had FastTree support values >0.9. Lineages are coloured and BAPS group designations are indicated. The scale bar indicates the number of SNPs represented by the horizontal branches. The location of ANZ045 *M. chimaera* reference genome is shown in green typeface. (B) Visualization using the Blast Ring Image Generator (32) of DNA:DNA whole genome comparisons among a subset of mycobacterial genomes used to identify *M. chimaera* specific regions. ANZ045 *M. chimaera* reference chromosome is depicted by the inner black circle. The identity of the subsequent rings is given in the legend. Annotations on the outer most ring show the location of the seven *M. chimaera*-specific regions, highlighting the region targeted for TaqMan PCR assay development.

**Fig. 2.**
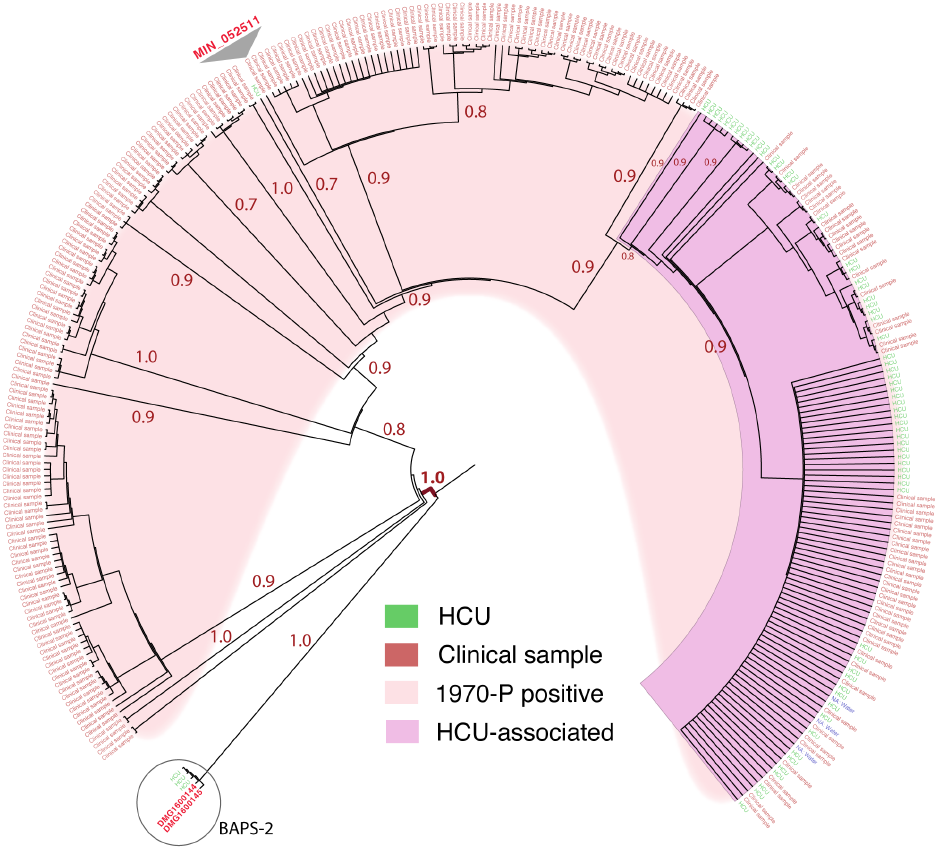
**Focused phylogenetic analysis of 303 *M. chimaera* genomes.** (A): Core genome maximum likelihood phylogeny of 303 *M. chimaera* (BAPS-3) and five other, related mycobacteria (BAPS-2), based on alignment of 10,166 variable nucleotide positions (recombination removed). The tree was inferred with FastTree using a GTR model of nucleotide substitution and rooted using DMG1600144 (BAPS-2, encircled) as an outgroup. The FastTree support values for major branches are indicated. Branch lengths have been transformed and they are proportional but not to scale. The location within the phylogeny of a sequence identified previously as *M. intracellulare* (MIN_052511) is indicated.

### In silico *genome comparisons to identify* M. chimaera *specific sequences*

A subset of 46 genome *M. chimaera* genome sequences as defined above became the ‘training set’ to find DNA segments present only in *M. chimaera* (Fig. 2, Table S1). Mapping of DNA sequence reads to the ANZ045 reference genome (refer methods) allowed the identification of 159 genomic segments >500 bp in length and covering 510,924 bp that were present across the 46 *M. chimaera* isolates and absent from eight *M. intracellulare* isolates (Fig. 1A). BLAST comparisons of the 159 segments against all entries in the NCBI Genbank *nt* database and removal of any non-*M. chimaera* specific sequence reduced the number to 37 segments (covering a total of 37,890 bps). A larger validation set comprising the 63 *M. chimaera* genomes described in Fig. 1A and 242 additional, publicly available *M. chimaera* genomes that satisfied our above phylogenomic inclusion criteria was then screened (Table). Six of the 37 *M. chimaera*-specific regions (SR), covering a total of 8,292 bp, were present in all 305 *M. chimaera* genomes (Fig. 1B). The six SRs ranged in length from 531 bp to 4,641 bp, the majority overlapping predicted chromosomal protein-coding sequences (Fig. 1B, Table 1). Inferred functions of these CDS are summarized (Table 1). The regions were scanned for sequence polymorphisms and one of these regions (SR1) that was 100% conserved among all *M. chimaera*, was selected as a template for the design of a TaqMan assay (Fig. 2, Table 2). SR1 spans two predicted CDS that DNA composition analysis and gene annotation predicted lay within a 35 - kb putative prophage or integrative mobile element. A 79 bp TaqMan amplicon (assay ID: 1970) was designed within a 2934 bp CDS (predicted function: unknown) (Table 2).

**Table 1.** Summary of the six putative *M. chimaera*-specific DNA segments identified by comparative genomics

| Region ID | Position in<br>ANZ045 chromosome | Length<br>(bp) | Putative CDS spanning region | TaqMan probe<br>ID<sup>1</sup> |
| --- | --- | --- | --- | --- |
| SR1 | 2047981-2052621 | 4,641 | 2 x hypothetical proteins | 1970 |
| SR2 | 2168889-2169420 | 531 | 1 x hypothetical protein |  |
| SR3 | 2171047-2171975 | 928 | 1 x hypothetical protein |  |
| SR4 | 2174560-2175316 | 756 | No protein-coding region detected |  |
| SR5 | 4855157-4855879 | 722 | 1 x hypothetical protein |  |
| SR6 | 5338898-5339612 | 714 | tRNA-His(gtg) |  |

Notes: <sup>1</sup>Probe ID number is based on the position of the first nucleotide of the TaqMan probe in the DNA segment (refer Table 2)

**Table 2.**
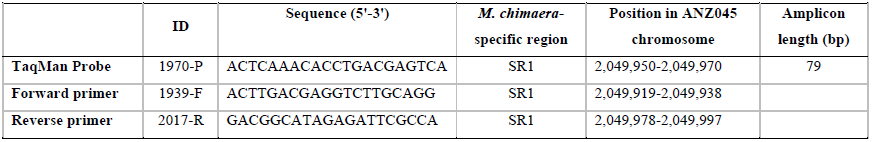
Sequences of the *M. chimaera*-specific TaqMan primers and probe

### TaqMan assay specificity testing

The above *in silico* analyses predicted the TaqMan assay would be diagnostic for the presence of *M. chimaera*. To test this prediction, DNA was prepared from 42 mycobacteria (including two *M. chimaera* isolates) and 22 other bacteria from 21 different genera. A pan-bacterial 16S rRNA PCR was first performed to ensure detectable bacterial DNA was present. All 64 DNA samples were positive by 16S rRNA PCR (data not shown) but only the two *M. chimaera* isolates were 1970-P TaqMan assay positive, supporting the *in silico* predictions that these assays are specific for *M. chimaera* (Table S2).

### TaqMan assay efficiency and sensitivity testing

To establish the limit of detection and amplification efficiency for the 1970-P TaqMan assay, 10-fold serial dilutions of purified *M. chimaera* ANZ045 genomic DNA were tested in triplicate. The 1970-P assay showed excellent performance characteristics, with a very good linear response across five orders of magnitude, R^2^ values >0.99, amplification efficiencies of 94%, and a detection limit around 20 genome equivalents (Fig. 3A). The detection limit was also assessed using dilutions of *M. chimaera* culture spiked into whole blood. Again, the assay showed excellent performance characteristics in this simulated clinical condition, with an absolute detection limit around 10 CFU (equivalent to 100 CFU/mL of blood). These experiments indicate 1970-P is a suitable qPCR assay for sensitive and quantitative detection of *M. chimaera* DNA.

**Fig. 3.**
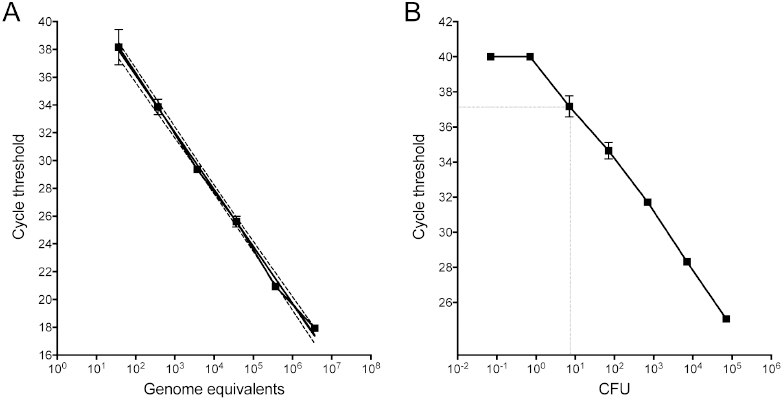
**Sensitivity testing of 1970-P TaqMan PCR for *M. chimaera*.** (A) Standard curve showing Ct versus *M. chimaera* 10-fold serial dilutions, expressed as genome equivalents (log scale). Shown are the mean and standard deviation for triplicate DNA preparations for assay 1970-P. A curve was interpolated using linear regression, R^2^= 0.992. Dotted lines indicate 95% confidence intervals. (B) Detection sensitivity of 1970-P for detection of *M. chimaera* spiked into whole blood, indicating a limit of detection of 10 CFU. Shown are the mean and standard deviation for triplicate blood samples and DNA extractions for each dilution.

### Detection of M. chimaera DNA in environmental samples

A key requirement for this assay is the ability to screen water and biofilm samples from heater-cooler units for *M. chimaera*, to determine if maintenance procedures have removed the bacteria from contaminated units, or prevented contamination. Having assessed the sensitivity and specificity of the assay with laboratory-prepared samples, we next explored performance with environmental samples. We screened concentrates from 33 water samples obtained from heater-cooler units at seven hospitals and HCU distributors in our region, that had been assessed by culture for *M. chimaera*. A total of 25 of 33 samples were positive by 1970-P TaqMan PCR, with estimated *M. chimaera* concentrations ranging from 2-102,000 GE/mL of water (Table 3). Using the culture results as a ‘gold standard’, the negative predictive value for the TaqMan assays was high (100%) with all eight culture positive samples also positive 1970-P TaqMan PCR (Table 4). However, there was poor correspondence between culture negative samples and qPCR. Fifteen samples negative by culture returned 1970-P TaqMan positive results, with Ct values for some of these samples less than 24, indicating high *M. chimaera* concentrations above 10,000 GE per milliliter of original sample (Table 3). Heterotrophic colony count (HCC) at 37ºC is used as a general indicator of water cleanliness, and in some settings may be used as a surrogate indicator of decontamination effectiveness (27). However, we observed a poor correlation between HCC and the presence of *M. chimaera* as measured by qPCR (Spearman’s ρ=0.2813, p=0.1128) (Table 3), suggesting that HCC is not a suitable surrogate for the presence or absence of *M. chimaera*.

**Table 3.** Environmental sampling qPCR, culture and heterotrophic plate count summary

| SAMPLE ID | 1970-P (Ct) | Genome equivalents | Volume water filtered (mL) (qPCR) | Genome equivalents/mL water | Culture positive (50 mL) | HCC/mL (log<sub>10</sub>) | Source |
| --- | --- | --- | --- | --- | --- | --- | --- |
| 17444 | No Ct | ND | 500 | ND | no | 2.7 | Facility C: HCU |
| 17651 | No Ct | ND | 200 | ND | no | 3.2 | Facility F: HCU |
| 17851 | No Ct | ND | 216 | ND | no | ND | Facility D: HCU cardioplegia tank |
| 18078 | No Ct | ND | 420 | ND | no | ND | Facility G: HCU cardioplegia tank |
| 18298 | No Ct | ND | 480 | ND | no | ND | Facility G: HCU cardioplegia tank |
| 18849 | No Ct | ND | 220 | ND | no | ND | Facility G: HCU cardioplegia tank |
| 18850 | No Ct | ND | 130 | ND | no | ND | Facility G: HCU cardioplegia tank |
| 19130 | No Ct | ND | 420 | ND | no | 1 | Facility D: HCU |
| 19330 | 37.76 | 40 | 1000 | 2 | no | ND | Facility G: HCU cardioplegia tank |
| 17443 | 37.67 | 42 | 34 | 62 | no | 2.8 | Facility C: HCU |
| 14211 | 36.17 | 98 | 270 | 18 | yes | ND | Facility C: HCU |
| 18079 | 36.04 | 105 | 960 | 5 | no | ND | Facility G: HCU cardioplegia tank |
| 17164 | 35.22 | 167 | 60 | 139 | no | 3.1 | Facility E: HCU |
| 9221 | 34.32 | 277 | 60 | 231 | yes | ND* | Facility A: HCU overflow water bottle |
| 18297 | 34.32 | 277 | 1000 | 14 | no | ND | Facility G: HCU cardioplegia tank |
| 18080 | 33.79 | 372 | 420 | 44 | no | 1 | Facility G: HCU cardioplegia tank |
| 10895 | 32.96 | 593 | 50 | 593 | yes | ND | Facility B: HCU unit-3 Patient circuit |
| 10896 | 32.65 | 706 | 70 | 504 | yes | ND | Facility B: HCU unit-3 cardioplegia circuit |
| 19128 | 32.53 | 755 | 420 | 90 | no | ND | Facility D: HCU |
| 10892 | 32.26 | 879 | 50 | 879 | yes | ND | Facility B: HCU unit-1 cardioplegia circuit |
| 10894 | 31.85 | 1106 | 30 | 1843 | yes | ND | Facility B: HCU unit-2 cardioplegia circuit |
| 18081 | 30.46 | 2412 | 960 | 126 | no | 1 | Facility G: HCU cardioplegia tank |
| 17850 | 30.19 | 2806 | 300 | 468 | no | 1 | Facility D: HCU cardioplegia tank |
| 19129 | 30.13 | 2902 | 400 | 363 | no | 1 | Facility D: HCU |
| 16569 | 29.19 | 4917 | 200 | 1229 | no | 4.3 | Facility D: HCU unit-1 |
| 18949 | 29.13 | 5086 | 220 | 1156 | no | ND | Facility H: HCU cardioplegia circuit |
| 18948 | 28.69 | 6509 | 275 | 1183 | no | ND | Facility H: HCU patient circuit |
| 18951 | 28.58 | 6923 | 280 | 1236 | no | ND | Facility H: HCU cardioplegia circuit |
| 10897 | 25.98 | 29765 | 70 | 21261 | yes | 1 | Facility B: HCU unit-1 patient circuit |
| 16571 | 24.82 | 57055 | 200 | 14264 | yes<sup>#</sup> | 1 | Facility D: HCU unit-3 |
| 10893 | 23.55 | 116325 | 80 | 72703 | yes | 2.7 | Facility B: HCU unit-2 Patient circuit |
| 18950 | 23.04 | 154851 | 130 | 59558 | no | ND | Facility H: HCU patient circuit |
| 16570 | 21.31 | 408655 | 200 | 102164 | yes | 1 | Facility D: HCU unit-2 |

Notes: <sup>#</sup>culture contaminated; \* ND = not detected

**Table 4.** Correspondence between M. chimaera culture and 1970-P qPCR in HCU samples

|  | Culture<br>positive | Culture<br>negative | Total |
| --- | --- | --- | --- |
| qPCR + | 10 | 15 | 25 |
| qPCR - | 0 | 8 | 8 |
| Total | 10 | 23 | 33 |

## Discussion

Since 2012 there have been a small but increasing number of case reports of invasive infection with *M. chimaera* in individuals who have undergone surgical procedures requiring cardiac bypass (8, 10-16). Almost all cases have involved placement of prosthetic valves or other prosthetic material and are linked to use of a specific type of heater cooler unit (HCU) in the bypass procedure (8, 13, 14). As contamination of these HCUs may have occurred at or near the time of manufacture and the machines are widely exported, exposure to *M. chimaera* during cardiac bypass surgery is an emerging issue in infection control that is not restricted by region or country (8). However, while it appears that generation of aerosols when machines are contaminated may be relatively common, so far the likelihood of infection for any exposed individual is very low (13). From a clinical perspective, this poses a major diagnostic challenge. Post-cardiac surgery *M. chimaera* infection has a long incubation period, non-specific symptoms and can be misdiagnosed as a steroid-requiring inflammatory condition with potentially disastrous consequences (10, 15). Moreover, there is a significant case fatality rate even when correctly identified and current opinion is that early accurate diagnosis will be key to achieving the best treatment outcomes (28). Given the non-specific symptoms and low prior probability of infection in a large exposed population, clinicians urgently need access to a specific and sensitive test for *M. chimaera* cardiac and extra-cardiac infection. Infection control practitioners have a different but equally challenging problem with respect to surveying and cleaning contaminated HCUs. In this report, we describe and validate a DNA target that is diagnostic for the presence of *M. chimaera*. We have shown under simulated conditions that it can accurately detect *M. chimaera* in human blood samples at very low concentrations and outperform culture in detecting *M. chimaera* in specimens obtained from contaminated HCUs.

The European Centre for Disease Prevention and Control recommends *M. chimaera* identification is performed by sequencing at least two conserved fragments among 16S–23S rRNA ITS, 16SrRNA, *rpoB* and *hsp65* [21,22]. Some laboratories have also used the MIN-2 probe in the INNO-LiPA Mycobacteria v2 line probe assay (6, 18). Here we simplify this suite of tests, with a *M. chimaera*-specific PCR assay, that has the advantages of providing a rapid yes/no result and an estimate of bacterial concentration. The test could be further enhanced with multiple DNA targets. The other five *M. chimaera*-specific regions reported here could be used to develop additional diagnostic targets (Fig. 1B). We envisage that this test will be used in conjunction with efforts to culture *M. chimaera* from specimens, as isolates are required for WGS to establish the genetic relatedness of isolates, and potentially for antimicrobial susceptibility testing (although antimicrobial resistance is not thought to be a major problem). As a previous risk assessment by Public Health England suggested a possible legionellosis risk for staff and patients, we propose that our assay should form part of a ‘HCU panel’ along with *Legionella* PCR (28, 29). While we have designed an assay to detect all *M. chimaera*, it may also be possible to detect specific *M. chimaera* lineages. For instance, it may be possible to use DNA deletion polymorphisms to discriminate among intra-species lineages, as in the *Mycobacterium tuberculosis* complex (30). We are currently exploring this possibility.

The infection risk posed by the water reservoirs within HCUs and the need for regular maintenance has been long recognized (29). Heterotrophic colony counts (HCC) are being considered as surrogates to assess the microbiological quality of HCUs (27, 31). We (like others) found a poor correlation between the presence of *M. chimaera* and HCC in water samples from HCUs (27), with examples of *M. chimaera* concentrations of 60,000 GE/mL when HCC were below the limit of detection (Table 3). More evaluation of the 1970-P assay is required, but our data suggest HCC may have an unacceptably high false-negative rate, which significantly reduces its utility for measuring the effectiveness of HCU decontamination procedures.

Screening HCU water samples with our TaqMan assay indicated the widespread presence of *M. chimaera* and a poor correlation with culture. There are several potential explanations for these observations. Despite the extensive *in silico* assessments, it is possible that the qPCR assay lacks specificity for *M. chimaera*, or that the culture method lacks sensitivity, or perhaps DNA from *M. chimaera* is still present but source organisms are no longer viable. Given the extensive *in silico* validation undertaken here to ensure target specificity, and the high prior probability that these HCU water samples contained *M. chimaera*, the discrepancy between culture and qPCR might best explained by either lower sensitivity of the mycobacterial culture method or PCR detection of intact DNA from non-viable *M. chimaera*. The later explanation is perhaps the most likely given that some of these PCR positive-culture negative samples were obtained from HCUs subjected to extensive decontamination procedures involving extended heating above 70ºC.

In summary, we have developed a new diagnostic tool for rapid, sensitive and specific detection of *M. chimaera* to help address the urgent need to screen patient and HCU samples.

## Acknowledgments.

We thank submitting health-care facilities and laboratories for providing water samples and mycobacterial isolates. We are grateful to Chris Coulter for provision of materials and critical review of the manuscript.

## Supplementary material

**Tables**:

Table S1: Mycobacterial genome sequences used in this study

Table S2: Summary of TaqMan assay in vitro specificity screening

